# A single-cell transcriptome analysis and prognostic model construction of stromal cells for renal cell carcinoma

**DOI:** 10.1101/2023.09.03.556072

**Authors:** Kuo Liao, Yifan Wang, Shuangxin Liu, Quhuan Li

## Abstract

Renal cell carcinoma (RCC) is among the top three cancers of the urinary system and its incidence keeps increasing worldwide in recent decades. However, methods for accurate prognosis evaluation and effective treatment are still lacking nowadays. Here, to explore the molecular expression features of RCC and establish a new RCC clinical prognosis evaluation model, a cell landscape of 187,263 renal cells obtained from eight patients with RCC was analyzed in this study. And by extracting and focusing on the main stromal cells from RCC tissues, innovative molecular characteristics and pathways of tumors were identified, like the well-known hypoxia pathway. By analyzing cell-cell communication, fibroblasts were found to promote tumor development by repressing natural killer cells. Based on Cox and least absolute shrinkage and selection operator regression analysis, four risk factors were screened and used to construct a reliable RCC clinical risk estimation model. In conclusion, our work provides new insights into the tumor microenvironment of RCC, as well as potential therapeutic targets and a clinical risk model for RCC invasiveness. Hopefully, these findings will be useful for cancer research and clinical treatment in future.

## Introduction

Renal cell carcinoma (RCC), more often known as renal cancer, is a heterogeneous malignancy originating from kidney tubular epithelial cells. Clear cell RCC (ccRCC) is the most common type of renal cancer. Currently, RCC is among the top three cancers of the urinary system, with a total patient number of 431,000 in 2020 (Sung et al. 2021). Despite the high mortality rate, methods for accurate prognosis evaluation and effective treatment are still lacking (Gill et al. 2018).

Targeted therapy and immunotherapy are currently the preferred treatment options for metastatic or unresectable renal cancer (Grimm et al. 2019; Iacovelli et al. 2022). Currently, targeted therapy for RCC has mainly focused on angiogenesis-related or intracellular signal transduction pathways, such as multi-target tyrosine kinase inhibitors (TKIs), mTOR inhibitors, and VEGF inhibitors such as sunitinib and pazopanib. However, most of these studies have failed to show effective outcomes. Among these, only a few drugs are still used as first-line therapy for metastatic ccRCC (Nabi et al. 2018). Even so, a considerable number of patients still develop resistance to these drugs after several years of adjuvant therapy and easily relapse. In contrast, immunotherapy, such as anti-PD-1 antibody therapy, has been reported to be more effective than targeted therapy with TKIs. The overall response rate for RCC is only 37–58% (Grimm et al. 2019; Kotecha et al. 2019). The reason for this low effective rate might be due to the compensation mechanism of the tumor microenvironment (TME); some signal pathways of tumor cells are inhibited, and others can still transduce signals, thus affecting the therapeutic effect (Bedognetti et al. 2019). A multi-target combination therapy strategy may be a way to improve the effectiveness of treatment, and this makes it necessary to understand the mechanism of tumor development.

In recent years, owing to advances in sequencing technology such as single-cell sequencing (scRNA-seq), our understanding of RCC has reached a new phase. Unlike traditional sequencing technology, scRNA-seq can further reflect the heterogeneity of tumors, which is an important characteristic of malignant tumors. Since the first report revealed the heterogeneity of renal tumors, many studies have also reported the heterogeneity of various subtypes of RCC (Chen et al. 2022; Gong et al. 2021; Young et al. 2018). A thorough understanding of RCC heterogeneity may have an important impact on cancer diagnosis and treatment. However, it is still difficult to accurately determine the clinical stages and effectively interfere with RCC deterioration using the current evaluation system. Therefore, the development of accurate prognostic evaluation and diagnosis of clinical patients based on scRNA-seq technology is necessary and urgent.

The TME is generally considered a complex component of immune and stromal cells (Joyce and Pollard 2009). Stromal cells, including tumor-associated macrophages (TAMs) (Mantovani et al. 2017), and cancer-associated fibroblasts (CAFs) (Bond et al. 2021), can interfere with immune function through tumor-stromal cell interaction (Nakasone et al. 2012); therefore, stromal cells can serve as potential therapeutic targets. In renal cancer studies, significant heterogeneity has been found in the TME of different RCC subtypes (Bond et al. 2021; Hakimi et al. 2019), which may facilitate tumor malignancy to different degrees through the interaction between stromal cells. Hence, focusing on stromal cells to explore new characteristics or possible targeting pathways will provide a new perspective for RCC treatment.

Therefore, we analyzed the single-cell RNA sequencing data of RCC (Obradovic et al. 2021). The stromal cells from the RCC tumor and paracancerous tissue of eight patients were extracted to obtain a comprehensive understanding of the molecular characteristics of RCC. We found that several new markers could be identified as potential targets for RCC therapy, and that hypoxia was still the most significant pathway for RCC. Furthermore, we built a risk assessment model for clinical diagnosis based on the differentially expressed genes (DEGs) and validated it with clinical data from the Cancer Genome Atlas (TCGA) dataset. This work may promote the clinical diagnosis of RCC patients and provide a new understanding of the disease.

## Materials and methods

### Data acquisition

The single-cell RNASeq count matrix of RCC patients was downloaded from Mendeley Data (https://data.mendeley.com/datasets/nc9bc8dn4m/1) (Obradovic et al. 2021). The expression matrix, along with the clinical information of the patients from the TCGA KIRC dataset (which contained 508 tumor samples after filtering), were obtained from the University of California Santa Cruz (UCSC) Xena (http://xena.ucsc.edu/) (Wang and Furey 2009). Moreover, the expression profiling dataset (GSE14762) was downloaded from the Gene Expression Omnibus (GEO) database of the National Center for Biotechnology Information (NCBI) https://www.ncbi.nlm.nih.gov/geo/) (Wang et al. 2009), which contained 10 RCC tumor bulk RNA sequencing (bulk RNA-seq) samples and 12 normal tissue cases. The dataset for model validation was obtained from the International Cancer Genome Consortium Data Portal (ICGC Data Portal) (https://dcc.icgc.org/), which contained two projects named KIRC-US and RECA-EU (Thomas et al. 2010).

### Single-cell data processing

We selected 30 samples in this study, which included 16 tumor samples and 14 paracancerous samples from eight ccRCC patients. The R package Seurat (version 4.1.1) (Hao et al. 2021) was used for basic quality control (QC). In the QC process, single cells with less than 1000 unique cell molecular identifiers (UMIs) or with more than 10% mitochondrion-derived UMI counts were considered low-quality cells and removed. Doublets, which were composed of more than two cells that were artificially introduced during the sequencing process, were removed; cells with more than 15000 UMIs were regarded as doublets. After QC, single-cell data were normalized and standardized using the R package Seurat with default parameters.

Batch effects among patients were eliminated using the R toolkit Harmony (version 0.1.0) (Korsunsky et al. 2019), which is a fast, sensitive, and accurate integration tool for single-cell data. The top 30 principal components along with the top 3,000 variable genes were used in this process. Unsupervised clustering was conducted using the FindClusters function (resolution = 0.3) of Seurat and visualized using uniform manifold approximation and projection (UMAP) along with non-linear dimension reduction (dim = 1:15). Finally, 187,263 single cells were filtered and divided into 21 main clusters for further analysis. The major cell types were annotated based on classic markers obtained from various published papers and divided into 10 main cell types (**Fig. 1A**).

**Figure 1.**
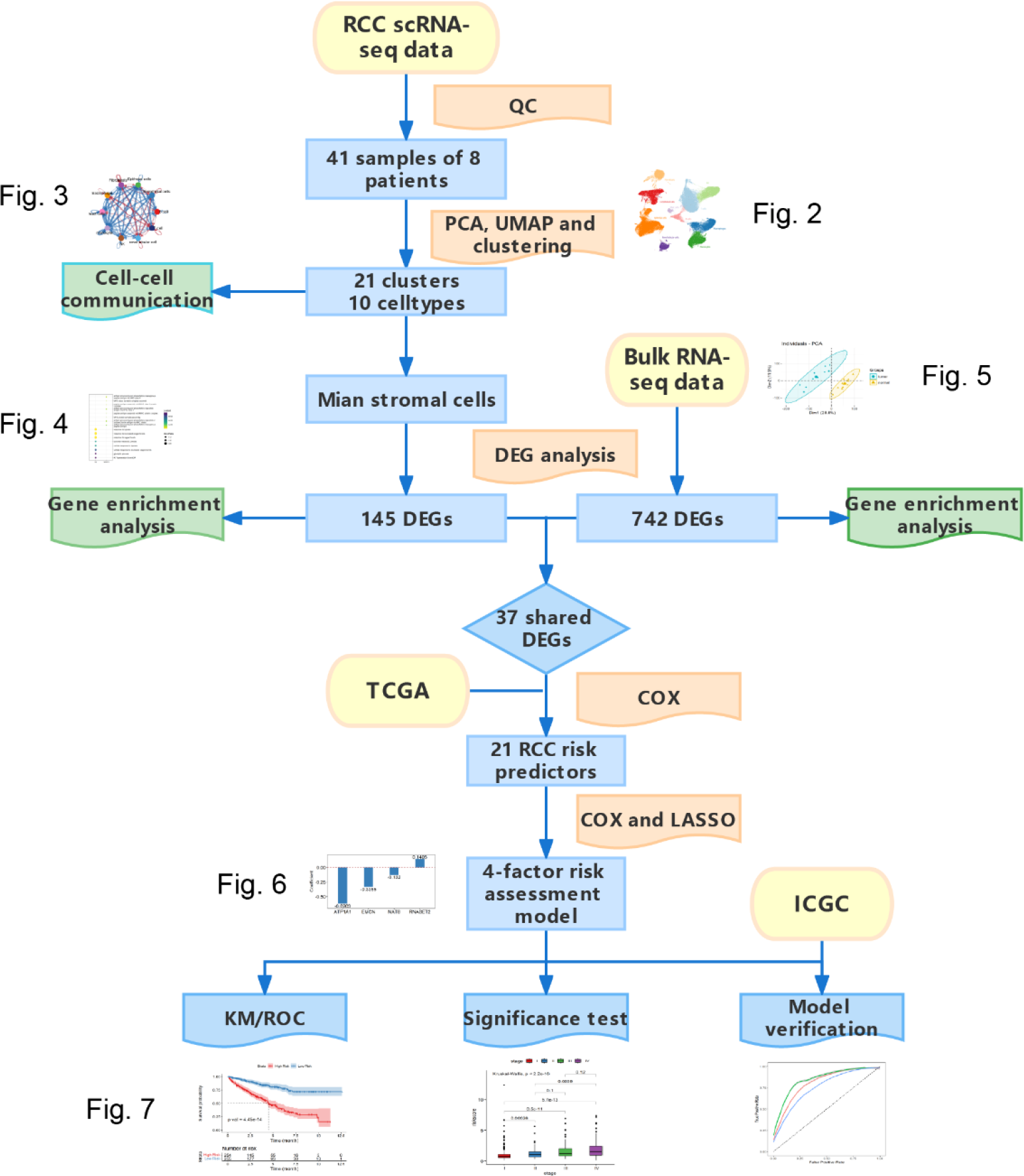
Flow chart

### Cell-cell communication analysis

A new R toolkit, CellChat (Jin et al. 2021) allows easier to explore the cell-cell communications for any given scRNA-seq dataset. The annotated RCC single-cell data were split into tumor and normal sample groups, and the interaction in all types of cells was analyzed using CellChat (version 1.1.3). The number and strength of ligand-receptor interactions were calculated and the key ligand-receptor pairs were recognized in tumor and normal samples; only those with a *p* < 0.05 were kept. Default parameters were kept in the CellChat analysis procedure, except for the dysfunctional signaling identification procedure using the differential expression analysis, where ligand.logFC was set as 0.5.

### Differential expression analysis

To focus on the variation in the main stromal cells in RCC, stromal cells were extracted from all samples. The three main stromal cell types in RCC single-cell data are epithelial cells, endothelial cells, and fibroblasts. Notably, because of the almost complete disappearance of renal tubular cells in the tumor tissues, they were not included in this differential analysis in the case of the over-weighted bias of the normal renal tubular cells. Due to technical deficiencies and the lack of gold standards, it is difficult to clearly distinguish and define true tumor cells and tumor-related cells in tumor tissues in terms of transcriptome data. Therefore, we assumed that all the stromal cells in the tumor tissue have synergy; thus, we focused on studying their systematization and holism. The DEGs of all the main stromal cell types between the tumor and paracancerous tissues were identified using the FindMarkers function of Seurat. The following cutoff threshold values (*p* < 0.001 and |log2FC| > 1) were used to filter the highly differentially expressed genes.

### Functional enrichment analysis

DEGs (*p* < 0.001 and |log2FC| > 1) after differential expression analysis were loaded into clusterProfiler (Yu et al. 2012) for Gene Ontology (GO) analysis in R. The pathways for which the adjusted p value was less than 0.05 were considered significantly enriched. The identified DEGs were used to perform gene set enrichment analysis (GSEA) using the R package clusterProfiler to detect significantly enriched pathways. Gene sets with false discovery rate (FDR) and *p* < 0.05 were considered significantly enriched. Here, the hallmark gene sets in MSigDB (https://www.gsea-msigdb.org/gsea/msigdb/collections.jsp) were chosen as enrichment targets.

### Bulk RNA-seq data analysis

Gene expression profile data (GSE14762) for renal cell carcinoma were downloaded from the GEO dataset, including 10 tumor samples and 12 paracancerous samples (Wang et al. 2009). The probe name in the data files was then changed to an international standard name. The limma package in Bioconductor (http://www.bioconductor.org/) was used for differential gene expression analysis of the bulk RNA sequencing data. After quantile normalization with cutoff setting |log2FC| > 1.5 and *p* < 0.001, 742 genes were screened and defined as highly differentially expressed genes. ClusterProfile in R was used for GO analysis, and only items with p < 0.05 were considered significantly enriched. (Fan et al. 2022)

### Establishment of a risk assessment model

The intersection of DEGs from the RCC scRNA data (|log2FC| > 1 and *p* < 0.001) and the bulk RNA-seq data (|log2FC| > 1.5 and *p* < 0.001) was loaded into univariate COX analysis with the R toolkit Survival. On the other hand, 508 RCC samples from TCGA were filtered out by a clinical follow-up time of more than 30 days, and the sample labeled ‘01A’ in the dataset was used for the univariate COX analysis. (Liu et al. 2022; Xiong et al. 2022)

Genes with *p* <0.001 in univariate COX analysis were retained and further subjected to LASSO-penalized COX analysis with lambda 100. Subsequently, five genes were retained and subjected to a multivariate COX analysis. Finally, a model with four risk factors was constructed and the corresponding risk coefficients were obtained. The risk scores of all TCGA samples were calculated using the predict.coxph function in the R package survival. Based on the obtained risk scores, all samples were divided into two dichotomized groups. Kaplan-Meier survival estimation was used to assess the accuracy of the model and show the survival difference between patients in the high- and low-risk groups, and the survivalROC package in R was used to calculate the receiver operating characteristic (ROC) curve of the model.

### Validation of the constructed risk model

Two projects (KIRC-US and RECA-EU) related to RCC were downloaded from the ICGC Data Portal and merged. Only samples with complete sequencing, clinical information, and a follow-up period of > 30 days were retained. A minimum of 488 samples were used for further validation. Normalized read counts in all samples were converted to fragments per kilobase million (FPKM) and subjected to log-normalization. The risk score of each RCC sample expression data from the ICGC database was calculated using the constructed risk model by the Survival in R with predict.coxph (type = ‘risk’). Then, Kaplan-Meier survival analysis and ROC curve were used to verify the model’s ability to distinguish high-risk from low-risk groups.

### Clinical evaluation by risk assessment model

By extracting the clinical information of RCC samples from all TCGA databases, grade (I–IV) and stage (NMT) well-defined samples were retained, and samples with unknown clinical information or vague stage information were dropped. The remaining samples were divided into two groups based on their estimated risk scores. A significance test was conducted between different clinical groups using the Kruskal-Wallis test.

## Results

### Cell landscape of the RCC and paracancerous tissues

Single-cell RNA-seq data from eight RCC patients, including CD45+ immune cells and CD45-other cells, were analyzed in our study. Through QC based on the expression levels of cell UMIs and mitochondrial genes, cells with average counts exceeding the filtration standard were filtered out, and 187,263 cells were finally obtained. Of these, 73,873 single cells were obtained from normal tissues, and the remaining were obtained from tumor tissues. After QC, Harmony (an R package) was applied to integrate all samples and to remove batch effects. In the integration process, the biological characteristics of CD45+ and CD45-cells were maintained for further study, which included the difference between the immune and stromal cells (**Fig. 2A and B**). After principal component analysis (PCA), dimensionality reduction by UMAP, K-nearest neighbor (KNN) self-clustering, and manual annotation by markers, these cells were divided into 10 major cell types: fibroblasts, epithelial cells, endothelial cells, renal tubular cells, macrophages, monocytes, mast cells, T cells, B cells, and NK cells (**Fig. 2A**). The expression levels of key markers reported in previous studies (Hu et al. 2020; Obradovic et al. 2021) for the 10 major cell types are shown in **Fig. 2C**. The proportion of cells in the RCC tumor and paracancerous tissues was then counted (**Fig. 2D**). Compared with the normal tissue, the immune cells, especially T cells, in the tumor showed a significant decrease, which was in accordance with the autoimmune decline that occurs in many tumors. The overall proportion of stromal cells increased, especially in epithelial cells and fibroblasts. Normal renal tubular cells disappeared completely in tumor tissues, which is consistent with previous studies (Chen et al. 2021; Hu et al. 2020; Sanchez and Simon 2018) and may be related to the transparency of RCC.

**Figure 2.**
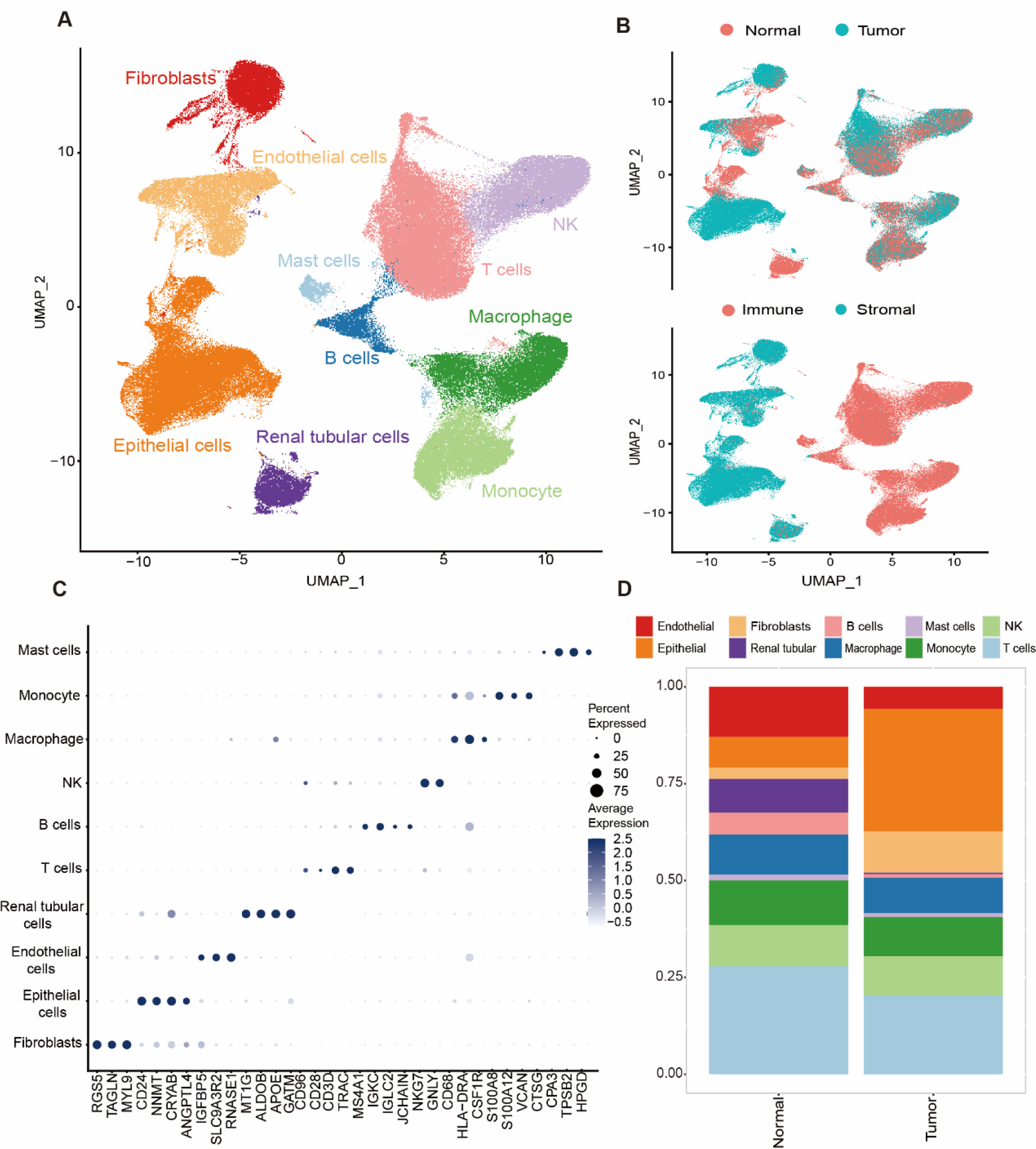
Overview of RCC single-cell data. **A**. UMAP plot of all the single cells of RCC, the 10 major cell types were labeled by different colors. **B**. UMAP plot of all the single cells samples originated from normal and tumor tissues, and UMAP plot of all the single immune (CD45+) or non-immune (CD45-) cells. **C**. Dot plot for the expression level of marker genes 10 major cell types. **D**. The fraction of cells originated from the 14 paracancerous and 16 tumor samples.

### Cell-cell communication difference between the RCC and paracancerous tissues

To explore the interaction between major cell types in the RCC microenvironment, we performed cell-cell communication analysis using the CellChat package in R. Compared with normal tissue, the number of cell interactions in tumor tissue decreased. However, the strength of cell interactions was significantly increased (**Fig. S1A and B**), which may be related to the degradation of differentiation function caused by mutations in stromal cells (Cairns et al. 2011). In addition, we found that monocytes showed the most significant changes in intercellular communication. Compared with paracancerous tissues, fibroblasts in the tumor showed the most significant increase in renal tubular cells (**Fig. 3A**). In terms of changes in cell interaction strength, epithelial cells showed the most significant enhancement of T cells in tumors (**Fig. 3B**). Specifically, compared with normal tissues, several receptor-ligand interactions related to tumor necrosis factors (TNFs) in RCC tumor tissues are remarkably activated, which is consistent with the effect found in many solid tumors (Balkwill 2009) (**Fig. 3C**). The significantly upregulated TNFs in RCC included CD70, CD40, and TWEAK. Among them, CD70, as the target of chimeric antigen receptor T-cell immunotherapy (CAR-T), has become a trend in renal cancer treatment (Drent et al. 2016). Therefore, it could be possible for CD40 and TWEAK, the other members of the TNFs family, to be new targets for CAR-T treatment in RCC. In addition, an increase in platelet-derived growth factor receptor (PDGF) was also noticeable (**Fig. 3C**). As an angiogenic factor, PDGF can induce the proliferation and migration of vascular endothelial cells, smooth muscle cells, and tumor cells, and further inhibit their apoptosis. These functions are closely related to tumor progression (Papadopoulos and Lennartsson 2018).

**Figure 3.**
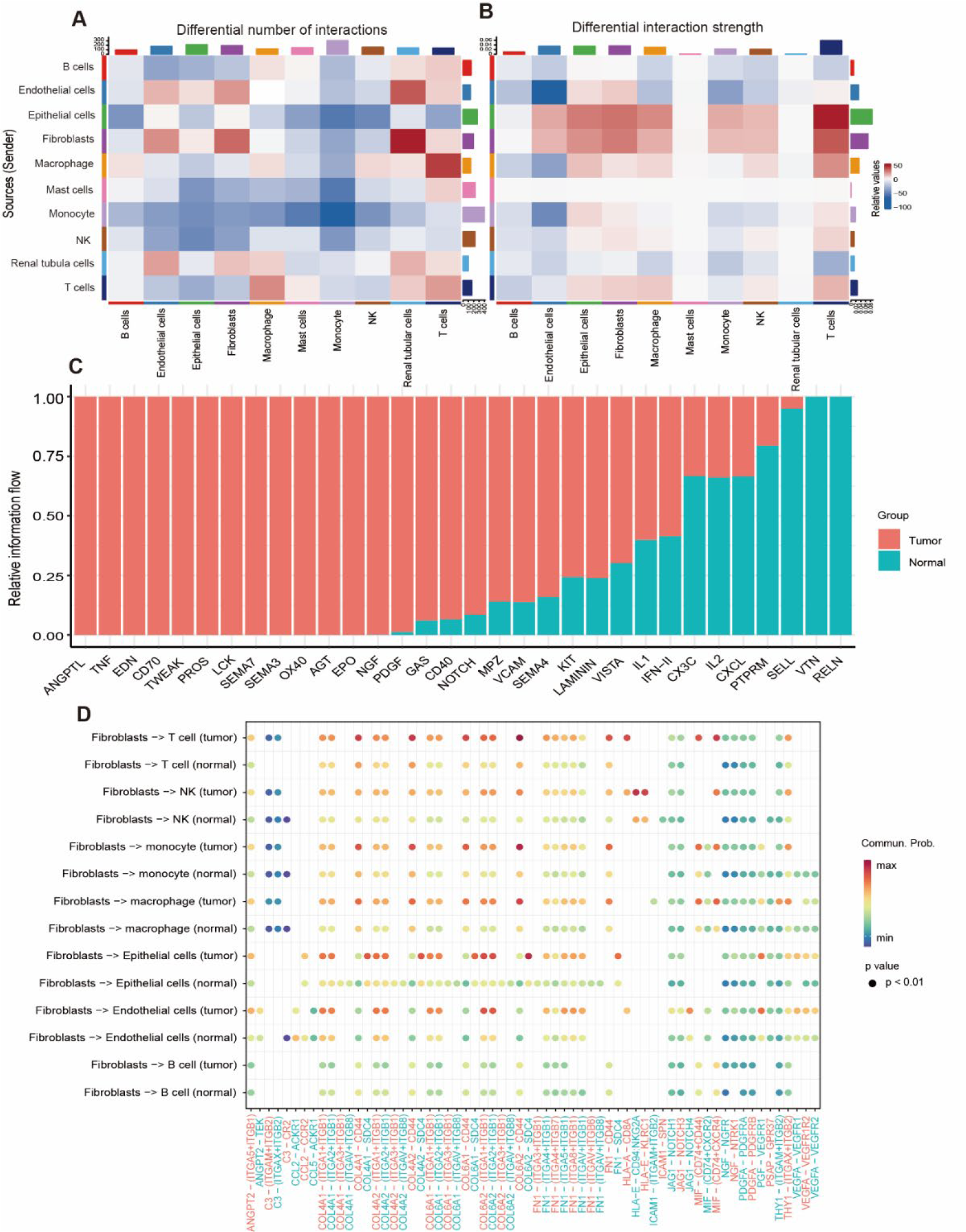
Cell-cell communication in RCC. (A, B) Heat map of the differential number of interactions (left) and interaction strength (right) in the cell-cell communication network between normal and tumor samples. Red and blue colors represent up-regulated and down-regulated signaling, respectively. (C) Bar plot of the significant signaling pathways. Genes were ranked by the relative information flow in the inferred networks of tumor (red) and normal (cyan) samples; *p* < 0.05 was considered as significant. (D) Bubble plot of the up-regulated signaling ligand-receptor pairs from fibroblasts to other main cell types in RCC.

For specific kinds of cell interaction, fibroblasts were picked as the sender to investigate the interactions with other receptor cells (**Fig. 3D**). As **Fig. 3D** showed, the communication probability of HLA-E-CD94:NKG2A pathway was highly improved between the fibroblast and NK cells in RCC. The NKG2A:CD94 complex is recognized by HLA-E, a major histocompatibility complex class I (MHC I) molecule. HLA-E is normally low-expressed, but it is upregulated in many tumors. Its expression induces the inhibition of cytokine secretion and the cytotoxicity of NK cells (Iwaszko and Bogunia-Kubik 2011), promoting tumor development. Signaling factors related to tumor metastasis such as COL6A1, COL6A2, COL4A1, and VEGFA were also highly expressed by the fibroblasts in RCC, with various targets (**Fig. 3D**). These interactions between the fibroblasts and other cell types in RCC further shed the light on the molecular mechanism between CAFs and the TME.

### Differential molecular characteristics and hypoxia-related pathways of RCC stromal cells

Considering the significance of stromal cells in the TME, we focused on the specific molecular characteristics of stromal cells in RCC. Since only a few normal renal tubular cells in the tumor regions were observed, we retained only three essential stromal cells in the differential gene analysis, including epithelial cells, endothelial cells, and fibroblasts. Using the Seurat package, we obtained 145 DEGs (|log2FC| > 1.0 and *p* < 0.001) from the main stromal cells between tumor and normal samples (**Fig. 4A**). Of these, 73 were upregulated and 72 were significantly downregulated. These DEGs also included some typical RCC markers reported in previous studies, such as NDUFA4L2, PLIN2, NNMT, CD70, CA9, and SLC17A3 (Dai and Sun 2020; Drent et al. 2016; Morrissey et al. 2015; Wang et al. 2017). Meanwhile, several innovative markers for RCC were first reported in this study, such as FABP7, REN, IGFBP3, CRYAB, WFDC2, and GAPDH (**Fig. 4A**). FABP7 was associated with fat synthesis, indicating a change in energy utilization in RCC (Kagawa et al. 2019). The CRYAB gene regulates molecular chaperones to maintain VEGF levels or promote tumor neovascularization, which has been reported in many tumors, such as breast cancer, lung cancer, colorectal cancer, and bladder cancer (Dai and Sun 2020). In addition, GAPDH showed hyperactivity in the mitochondria of RCC stromal cells. The upregulation of GAPDH reflected the large amount of energy required by tumor tissue and further implied the prominence of anaerobic respiration in RCC stroma. Notably, no significant change in the expression of these genes was observed in the immune cells (**Fig. S2**), further illustrating the uniqueness of the transcriptome heterogeneity in the stromal cell population.

**Figure 4.**
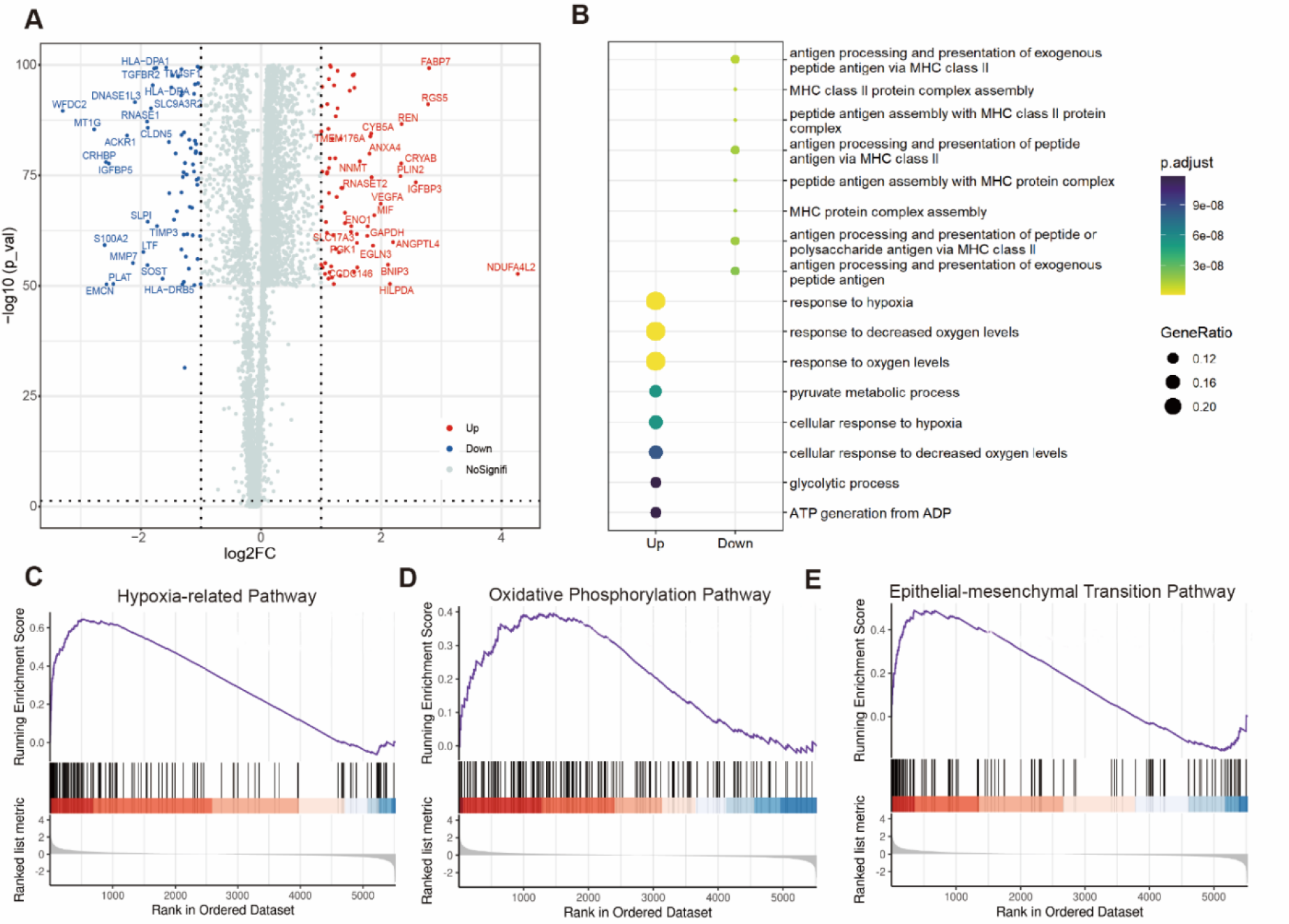
Differential expression genes and gene set enrichment analysis (GSEA). **A**. Volcano plot of differentially expressed genes (DEGs) of stromal cells between RCC and paracarcinoma tissue. The stromal cells include epithelial cells, endothelial cells, and fibroblasts. Upregulated genes were colored in red, and downregulated genes were colored in blue, while grey means no significant. Part of upregulated and downregulated genes were annotated, respectively. **B**. Dot plot of Gene Ontology analysis of DEGs. Upregulated and downregulated pathways significantly enriched were plotted (FDR < 0.05) . **C-E**. GSEA of stromal cells. The hypoxia-related pathway, oxidative phosphorylation pathway and epithelial-mesenchymal transition pathway were significantly enriched.

To better understand the biological changes of RCC stromal cells, we performed GO analysis for 145 DEGs. The results showed that the hypoxia-related pathway in RCC was highly enriched (**Fig. 4B**). Generally, hypoxia was one of the hallmark events of solid tumors. However, for RCC, we found this phenotype seemed to be particularly pronounced. This feature of RCC may suggest that targeting hypoxia-related factors may be an effective strategy to control the occurrence and development of RCC. When we took all of the weak DEGs (*p* < 0.05) into account by GSEA analysis, the results showed that the DEGs of RCC were still highly enriched in the hypoxia-related pathway (**Fig. 4C**). Besides, the oxidative phosphorylation pathway in RCC was also significantly enriched in RCC stroma (**Fig. 4D**), which may be related to aerobic glycolysis occurring in tumor stromal cells (Reinfeld et al. 2022). This high oxygen consumption in the low of oxygen supply environment could lead to further deterioration of the TME. Meanwhile, we found the epithelial-mesenchymal transition (EMT) pathway in RCC was also highly activated via GSEA (**Fig. 4E**). Although in this study, we didn’t force to separate tumor cells from epithelial cells in the tumor tissues, the high enrichment of MET process was in accord with our expectation. This could further confirm that RCC was originated from the epithelial cells, as we mentioned in the introduction.

### Key differential molecular characteristics of RCC tissue analyzed for bulk RNA-seq data

To further evaluate differential expression in RCC, we selected 22 RCC samples from GEO data, including 10 RCC tumor and 12 normal tissue samples, as objects for further analysis (**Fig. 5A**). Using gene differential expression analysis by limma, 704 significant DEGs between normal and tumor samples (|log2FC| > 1.5 and *p* < 0.001) were found. Therefore, 37 genes were intersected with the DEGs found in the RCC scRNA-seq data (**Fig. 5B**), including ATP1A1, ANXA4, TGFBI, HISTIHIC, HLA-DPA1 and DPA1. Based on the DEGs obtained from the bulk RNA-seq data, we also conducted GO analysis and found pathways similar to those from single-cell data. The response to hypoxia and oxygen level pathways in RCC was significantly increased (**Fig. 5D**), further confirming the role of hypoxia in RCC tumor cells. An improvement in the immune response related to phagocytosis and leukocyte activation has also been observed, which is consistent with previous findings in many solid tumors (Herrera-Campos et al. 2022; Terrén et al. 2019). The change in the prominent position from hypoxia- and oxygen-sensing pathways in scRNA-seq to immune response-related pathways in bulk RNA was mainly due to the differences in the analyzed objects. Only stromal cells were selected to study DEGs and biological pathways in the scRNA-seq data. Most importantly, although the DEGs showed some differences between bulk RNA-seq data and scRNA-seq data, there were some parts that confirmed each other, especially for the enriched pathways. This not only verified the accuracy of our single-cell analysis but also further highlighted the advantages of single-cell omics.

**Figure 5.**
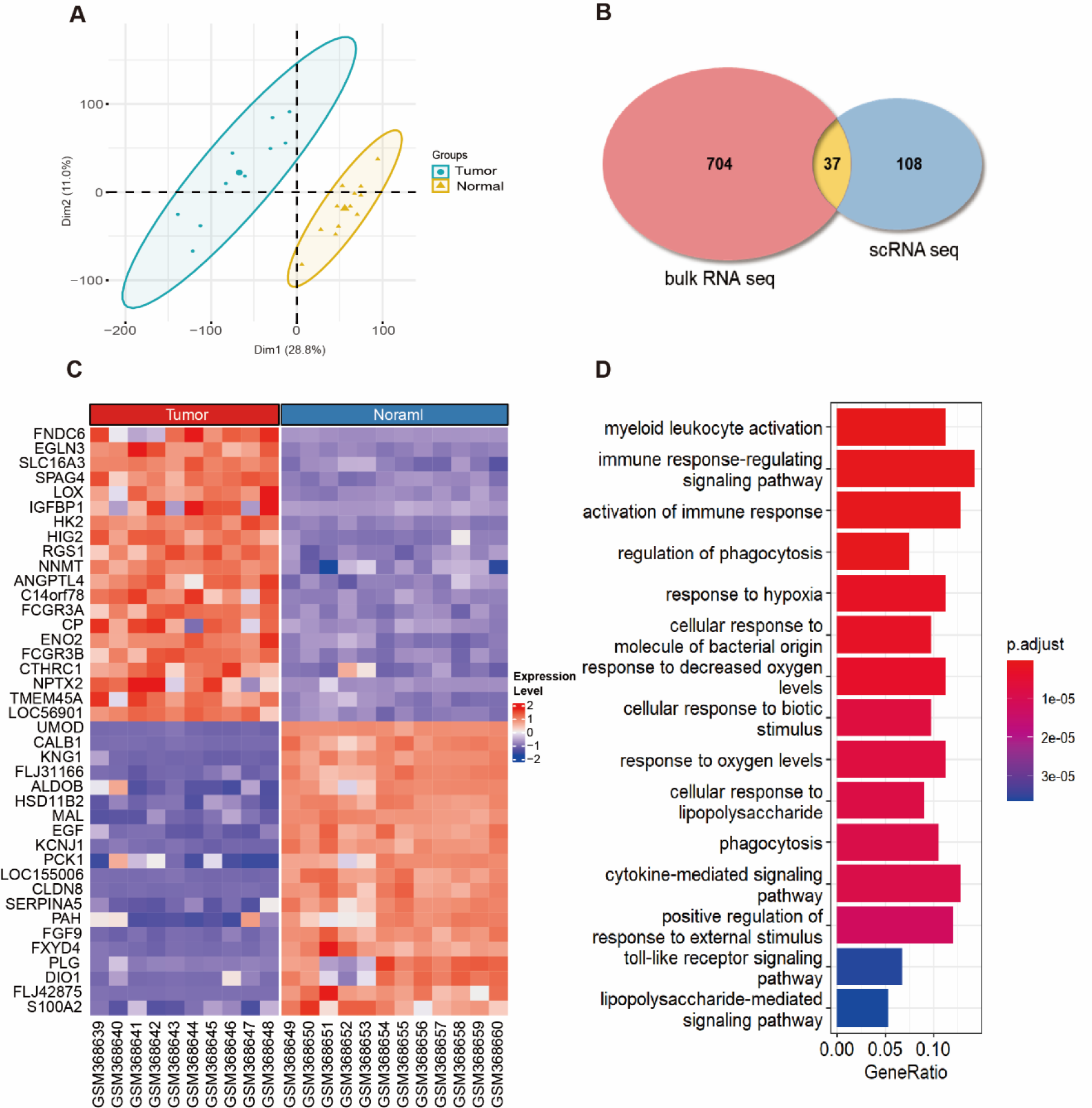
Molecular characteristics of RCC analyzed by bulk RNA-seq data. **A**. Two-dimensional principal component analysis (PCA). Blue circles and yellow triangles represented 10 RCC tumor samples 12 normal samples, respectively. **B**. Venn plot of the intersections of the differentially expressed genes (DEGs). The DEGs in blue oval were from stromal cells of scRNA-seq data analyzed by Seurat (*p* < 0.001 and |log2FC| > 1.0), and the DEGs in the red oval were from RCC tissue of GEO bulk RNA-seq data analyzed by limma (*p* < 0.001 and |log2FC| > 1.5). **C**. Heat map of the top 40 DEGs of bulk RNA-seq data, the column names are samples names. **D**. Bar plot of Gene Ontology pathway analysis for bulk RNA-seq DEGs. Only the top 15 pathways were plotted. FDR < 0.05 was considered as significantly enriched.

### Establishment of a risk assessment model and finding of potential targets for RCC clinical diagnosis and therapy

After differential expression analysis, 37 common DEGs (**Fig. 5B and Table S1**) were identified for further screening to build a prognostic risk model for RCC. Through univariate COX regression analysis, 21 genes showed a strong correlation with the lifespan of RCC patients (*p* < 0.05) using data from TCGA, including EMCN, AQP1, and DNASE1L3, which can be potential biomarkers for the prognosis of RCC (**Fig. 6A**). We further conducted a LASSO regression analysis for the 21 candidate markers (*p* < 0.05) obtained from the univariate COX regression analysis. An RCC risk evaluation model was constructed with four key factors: ATP1A1, RNASET2, NAT8, and EMCN (**Fig. 6B**). Based on the model built in this study, 509 patients were assigned a risk score. Patients with higher scores tended to have lower survival times in both dead and live patients, whose survival times were counted until the end of follow-up (**Fig. 6C**). To further clarify the relationship between risk scores and patient survival probability, patients were divided into two groups based on their risk scores. The low- and high-risk groups included those with risk scores below and above the median risk. The Kaplan– Meier survival curve showed that the survival time of the low-risk group was significantly longer than that of the high-risk group (**Fig. 6D**). The discrimination (*p* = 5x10^-16^) of our analysis model based on stromal cells was high. After that, we conducted receiver operating characteristic (ROC) curve analysis for 5-year, 10-year, and 15-year clinic follow-up data of

**Figure 6.**
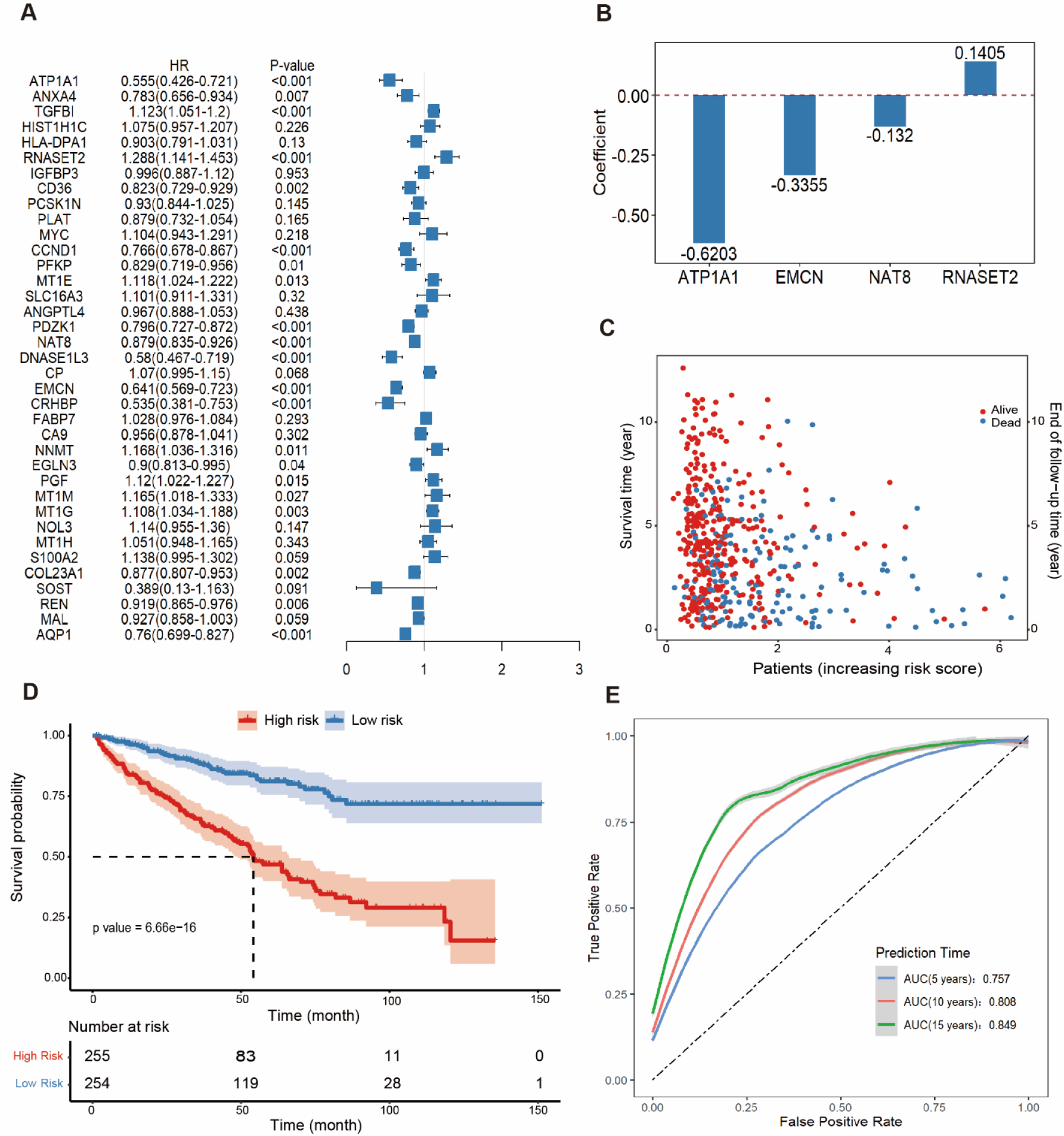
Development of the risk assessment model for RCC. **A**. The univariate COX regression analysis. HRs (hazard ratios) for 21 DEGs were obtained based on the correlation of genes with clinical patients survival from TCGA KIRC cohort. **B**. The construction of four-factors risk assessment model with multivariate COX regression analysis. It showed the coefficients of four factors. **C**. The distribution plots of risk scores. The risk scores were calculated with risk assessment model built in this study. The blue and red dots were survival times for dead patients and the end of the follow-up time for alive patients, respectively. **D**. Kaplan-Meier survival curve for patients collected in TCGA KIRC. Based on the risk scores calculated by the COX model, patients were divided into two groups. The low and high risk groups were those whose risk scores were below and above the risk median, respectively. Hazard ratios (HRs), with their 95% confidence intervals in brackets, were shown. **E**. ROC curve and AUC value of risk assessment model. The blue, orange, and green lines were the ROC curves for patiens with 5-year (AUC = 0.757), 10-year (AUC = 0.808) and 15-year (AUC = 0.849) follow-up times, respectively.

RCC patients from TCGA database, and finally obtained an ideal model evaluation index AUC value (maximum AUC was 0.849). This result proved the feasibility and accuracy of our risk assessment model for the prognosis of patients (**Fig. 6E**), which is based on the DEGs of tumor stromal cells. Compared with many traditional prediction models and some previously reported risk models (**Table S2**), our model has an advantage in the clinical evaluation of the prognosis of RCC patients with high accuracy and few required factors.

### Model validation and clinical evaluation with external data

Owing to the important role of stromal cells in the TME, immune cell infiltration, and RCC tumor metastasis, a series of significance tests was performed to investigate the relationship between the clinicopathological features of RCC and our risk model. The results showed that the risk scores were positively correlated with clinical stages (Kruskal-Wallis test, *p* < 0.001, **Fig. 7A**), especially when evaluating the degree of clinical in situ tumor deterioration of I/II and III/IV (**Fig. 7A**). In addition, there was a significant correlation between risk values and different grades of metastasis (Kruskal-Wallis test, *p* < 0.001, **Fig. 7B**). In addition, we conducted a risk assessment using the COX risk model for the external data obtained from the ICGC database, which contained the filtered transcriptional group expression data and clinical data of RCC patients. Kaplan-Meier survival analysis showed that our model was highly differentiated between the two groups of patients with relatively different risks. The high-risk group had a worse prognosis and shorter survival time than the low-risk group (**Fig. 7C**). Finally, ROC curves were constructed for a 5-year, 10-year, and 15-year prognostic evaluation of RCC patients, the AUC value indicated the reliability of our model (**Fig. 7D**).

**Figure 7.**
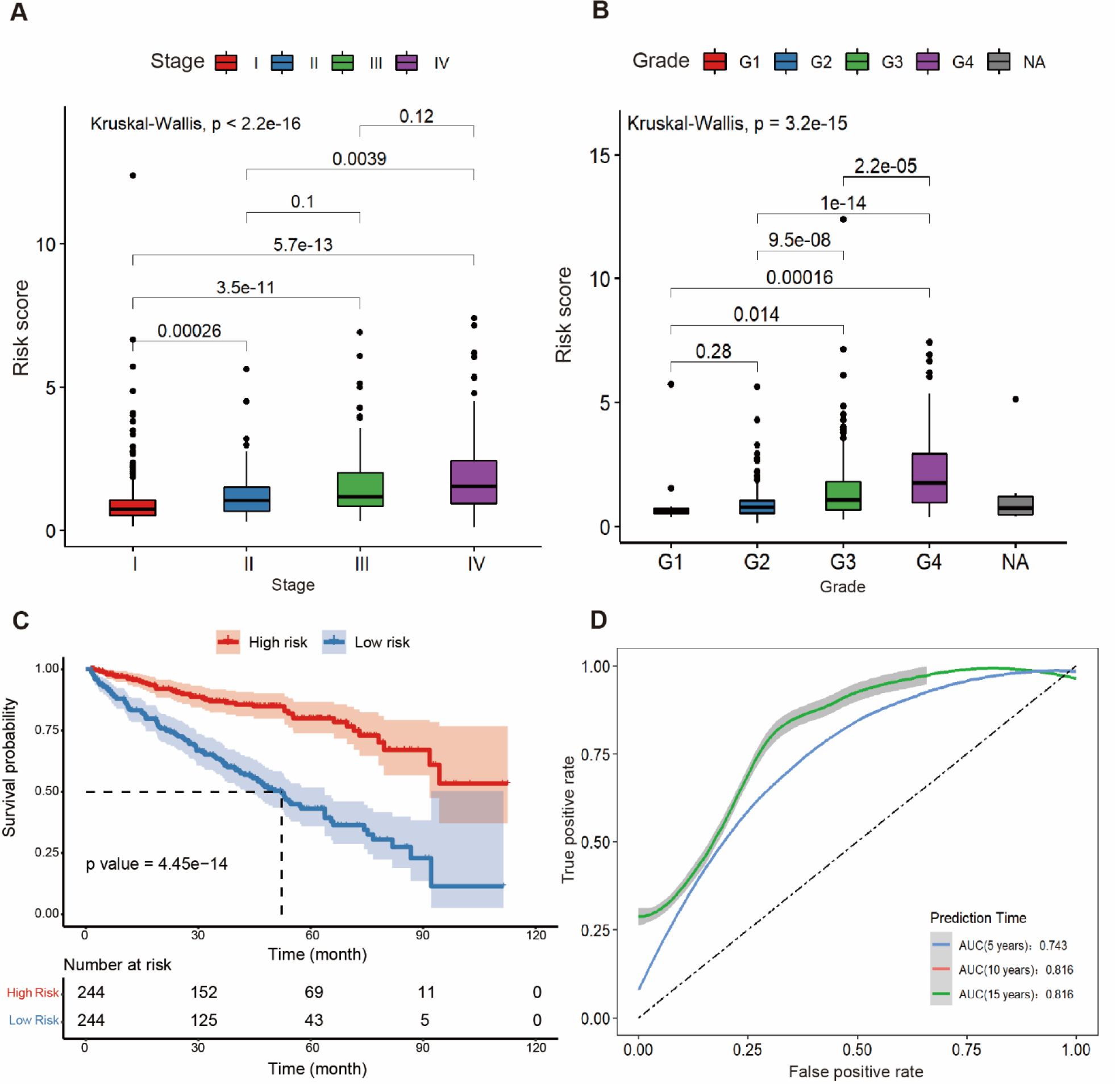
Estimation and verification of the risk model. **A**. The box plot of risk score with clinic stages (I∼IV). The Kruskal-Wallis test showed the significant difference (*p* < 2.2x10^-16^). **B**. The box plot of risk score with clinic grades (G1∼G4). The Kruskal-Wallis showed a significant difference (*p* < 3.2x10^-15^), and the NA refers to samples in TCGA without a precise grade. **C**. Kaplan-Meier survival curve for RCC patients collected from ICGC database. The risk scores were calculated with the COX model built in this study. **D**. ROC curves of risk model in predicting 5-year (in blue, AUC = 0.743), 10-year (in orange, AUC = 0.816) and 15-year (in green, AUC = 0.816) progression outcomes. Note, the curve of 10 years is overlapped by the one of 15 years.

## Discussion

In recent years, the incidence of RCC has rapidly increased in most countries and regions (Sung et al. 2021). However, a reliable prognostic evaluation method for clinical RCC is lacking. Although several biomarkers in RCCs have been identified, most of them cannot provide precise prognostic outcomes (Gill et al. 2018; Kotecha et al. 2019). Therefore, a multidimensional and precise clinical appraisal system is required for an efficient treatment.

In this study, we focused on RCC stromal cells to explore new markers and possible target pathways. Through the extraction of stromal cells from scRNA-seq data of tumor and para-cancerous tissues, we analyzed the most significant DEGs and revealed active pathways in stromal cells, such as hypoxia and oxygen sensing pathways. This was also supported by bulk RNA sequencing. Based on the differential expression analysis, EGLN3 and SLC16A3 may be upstream or promoting factors related to hypoxic deterioration. These findings provide a new basis for the clinical diagnosis and evaluation of RCC and further development of new RCC therapeutic drugs.

For a better understanding of cell interactions, we used the R package CellChat to conduct cell-cell communication analysis. The results showed that the number of cell interactions in the tumor tissue of RCC was reduced, but the strength of interaction was significantly increased, and some pathways were significantly activated, such as TNF and angiogenic factors. Moreover, fibroblasts were found to have a significant effect on NK cells, indicating that they were able to inhibit the immune effects of NK cells by releasing HLA-E.

Furthermore, we performed a single factor COX regression analysis of DEGs for the data obtained from TCGA database and found multiple factors that could be independently correlated with RCC prognosis, such as TIMP3 and PGK1. Then, LASSO regression and multivariate COX regression analyses, in combination with clinical follow-up data, were performed to construct an RCC prognostic risk prediction model. The excellent prediction performance was verified using data from TCGA and ICGC databases. Meanwhile, compared with various methods, this model has great discrimination ability for the different stages and grades of RCC, based on the molecular differences of stromal cells. Therefore, the model built in this study can provide a higher prediction accuracy with limited prediction factors compared with many existing models (**Table S2**) (Liu et al. 2021; Lv et al. 2021; Shi et al. 2017; Tusong et al. 2017; Zhang et al. 2021a; Zhang et al. 2021b).

In conclusion, the study of tumor stromal cells provides a new perspective for investigating the importance of molecules in the immune microenvironment of RCC. Several new tumor marker genes have been proposed as potential targets for the treatment of RCC. The establishment of an RCC clinical risk estimation model will also aid in clinical diagnosis and evaluation, thus further guiding the selection of treatment options. Although the effectiveness of this model was proven by clinical data from TCGA and ICGC databases, there is still much work to be done before effective drug development and RCC cure can be achieved. Whether the drugs that target stromal cells can effectively interfere with the metastasis and deterioration of the tumor needs to be verified through further experimental exploration in the follow-up work.

## Acknowledgements

We are very grateful to all these provided us with advice and guidance during this work.

## Author contributions

All authors contributed to the study conception and design. Material preparation, data collection and analysis were performed by Kuo Liao and Yifan Wang. The first draft of the manuscript was written by Kuo Liao and reviewed by Quhuan Li. All authors contributed to the article and approved the submitted version.

## Supplementary data

**Figure S1.**
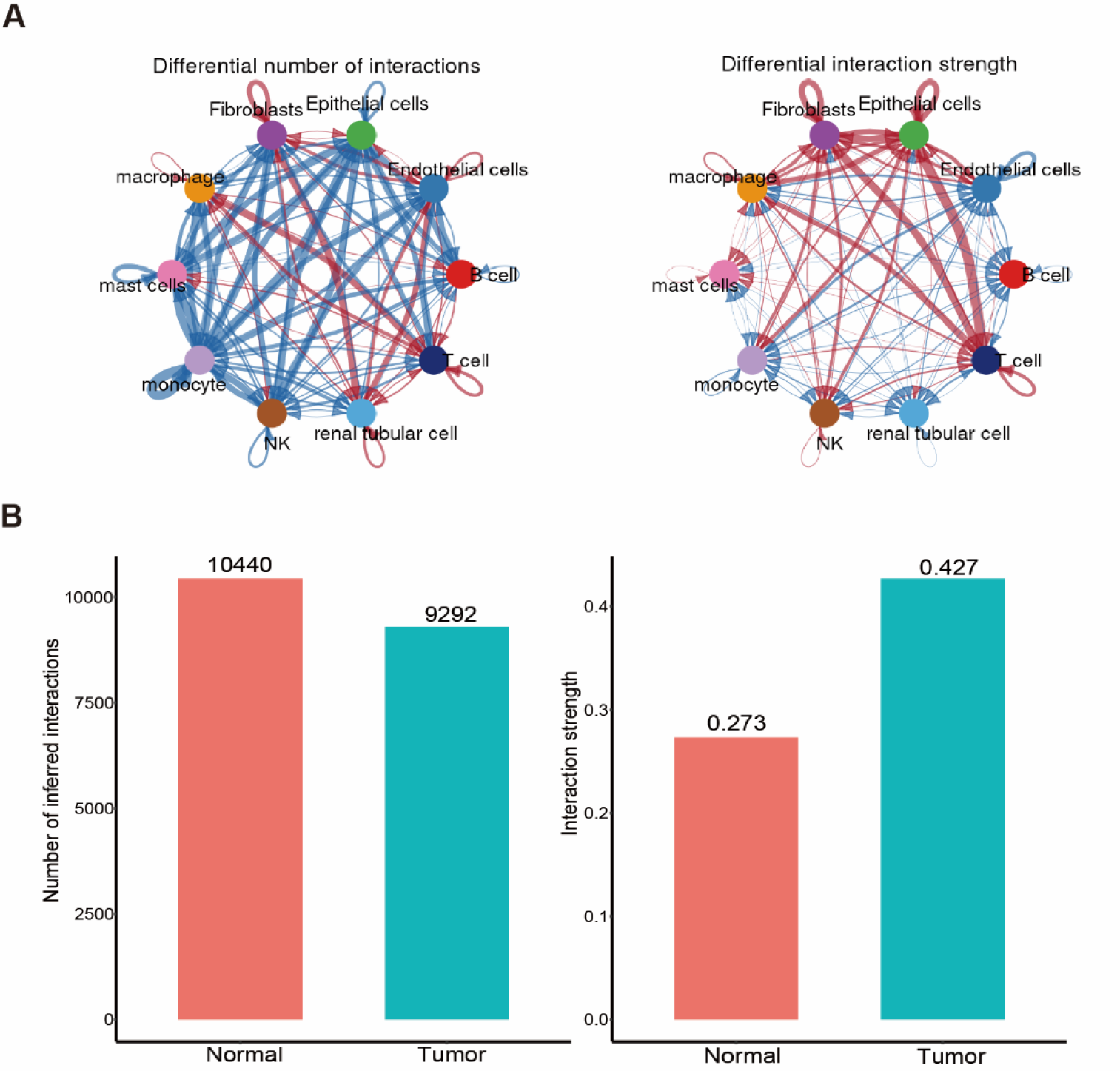
An overview of the number and strength of cell-cell communication. A) the pie chart for differential number and strength of interactions (blue for tumor tissue and red for paracancerous tissue). B) the bar chart for quantitative interactions.

**Figure S2.**
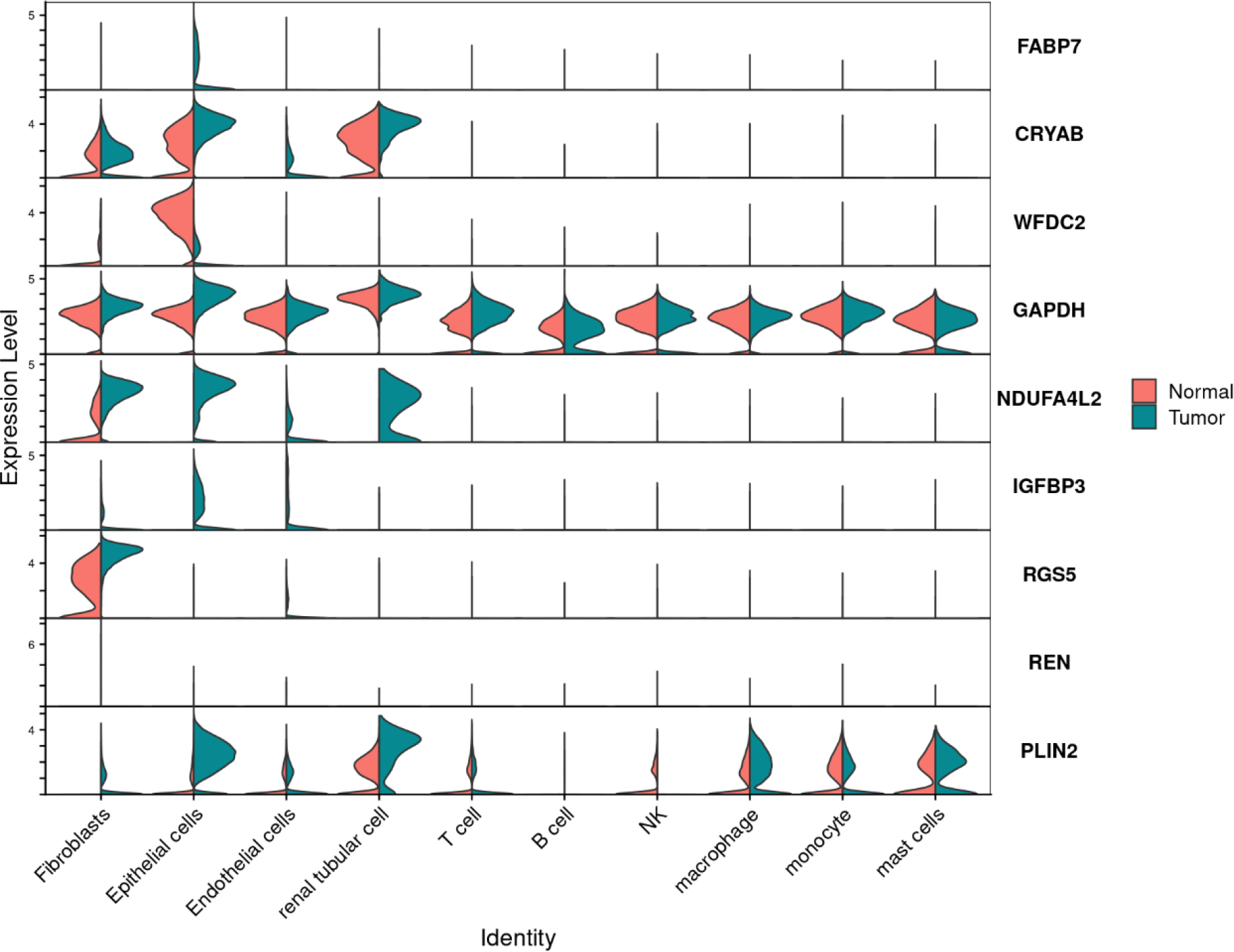
Violin plot of differentially expressed gene expression. The expression showed significant difference between tumor and normal tissues in the RCC stromal cells but not obvious difference in the immune cells.

**Table S1.** Results of the univariate COX regression of 37 DEGs.

| Gene | Coefficient | HR | Lower.95 | Upper.95 | p value |
| --- | --- | --- | --- | --- | --- |
| ATP1A1 | -0.58959 | 0.554556 | 0.426255 | 0.721474 | 1.12E-05 |
| ANXA4 | -0.24487 | 0.782806 | 0.655811 | 0.934394 | 0.006701 |
| **TGFBI \*** | 0.115926 | 1.122913 | 1.050578 | 1.200227 | 0.000644 |
| HIST1H1C | 0.071865 | 1.07451 | 0.956513 | 1.207062 | 0.225953 |
| HLA-DPA1 | -0.10214 | 0.902907 | 0.791063 | 1.030563 | 0.130084 |
| **RNASET2 \*** | 0.252807 | 1.287635 | 1.141323 | 1.452704 | 3.99E-05 |
| IGFBP3 | -0.00353 | 0.996471 | 0.886646 | 1.119901 | 0.952691 |
| CD36 | -0.19491 | 0.822912 | 0.729126 | 0.928763 | 0.001594 |
| PCSK1N | -0.07247 | 0.930093 | 0.843756 | 1.025263 | 0.144836 |
| PLAT | -0.12919 | 0.878806 | 0.732423 | 1.054444 | 0.164620 |
| MYC | 0.09868 | 1.103713 | 0.943271 | 1.291445 | 0.218222 |
| CCND1 | -0.26596 | 0.766468 | 0.677749 | 0.866801 | 2.26E-05 |
| PFKP | -0.18784 | 0.828748 | 0.718794 | 0.955522 | 0.009697 |
| MT1E | 0.111817 | 1.118308 | 1.023639 | 1.221731 | 0.013223 |
| SLC16A3 | 0.09613 | 1.100902 | 0.910797 | 1.330686 | 0.320264 |
| ANGPTL4 | -0.03377 | 0.966797 | 0.887791 | 1.052835 | 0.437579 |
| PDZK1 | -0.22771 | 0.796359 | 0.727494 | 0.871743 | 8.04E-07 |
| **NAT8 \*** | -0.12865 | 0.879281 | 0.835098 | 0.925801 | 1.00E-06 |
| DNASE1L3 | -0.54538 | 0.579624 | 0.467266 | 0.719 | 7.03E-07 |
| CP | 0.06737 | 1.069691 | 0.99514 | 1.149827 | 0.067582 |
| **EMCN \*** | -0.44421 | 0.641331 | 0.56856 | 0.723416 | 4.87E-13 |
| CRHBP | -0.62473 | 0.535408 | 0.380553 | 0.753278 | 0.000335 |
| FABP7 | 0.028071 | 1.028468 | 0.976005 | 1.083751 | 0.293352 |
| CA9 | -0.04491 | 0.956085 | 0.877973 | 1.041146 | 0.301736 |
| NNMT | 0.154983 | 1.167638 | 1.036372 | 1.31553 | 0.010861 |
| EGLN3 | -0.10585 | 0.899561 | 0.813314 | 0.994953 | 0.039556 |
| PGF | 0.113236 | 1.119897 | 1.022407 | 1.226682 | 0.014815 |
| MT1M | 0.152437 | 1.164669 | 1.017642 | 1.332938 | 0.026831 |
| MT1G | 0.102852 | 1.108328 | 1.034422 | 1.187514 | 0.003487 |
| NOL3 | 0.130787 | 1.139725 | 0.955038 | 1.360128 | 0.147072 |
| MT1H | 0.049658 | 1.050912 | 0.948353 | 1.164562 | 0.343220 |
| S100A2 | 0.129672 | 1.138455 | 0.995162 | 1.302381 | 0.058850 |
| COL23A1 | -0.13128 | 0.876968 | 0.806805 | 0.953232 | 0.002030 |
| SOST | -0.94319 | 0.389383 | 0.130371 | 1.162982 | 0.091122 |
| REN | -0.08465 | 0.918831 | 0.864907 | 0.976118 | 0.006082 |
| MAL | -0.07542 | 0.927357 | 0.8576 | 1.002787 | 0.058730 |
| AQP1 | -0.27391 | 0.760402 | 0.698985 | 0.827217 | 1.84E-10 |

Notes: The coefficient represented the correlation coefficient between the gene expression and clinic risk. The Hazard Rate (HR) represented the correlation of the gene expression with clinic risks, which >1 means risk factors and <1 means protective factors. The genes marked with star were the important indicators which were used to build the risk model.

**Table S2.** The establishment of several risk models for RCC.

| Indicator | Data source | AUC | p value | Ref. |
| --- | --- | --- | --- | --- |
| ATP1A1, EMCN, NAT8, and RNASET2 | Single-cell & bulk transcriptome | 0.757(5-year)<br>0.808(10-year)<br>0.849(15-year) | <0.0001 | This work |
| Seven glycolysis-related signatures (GALM, TGFA, RBCK1, CD44, HK3, KIF20A and IDUA) | Bulk transcriptome | 0.767 (5-year) | - | Lv et.al. 2021 |
| GTSE1, CENPF, SMC2 and H2AFV | Single-cell & bulk transcriptome | 0.712(3-year)<br>0.726(5-year) | - | Zhang et.al. 2021 |
| CAF-related signature (COL16A1) | Single-cell transcriptome | 0.6255 | <0.0001 | Liu et al. 2021 |
| CD248 | Bulk transcriptome | 0.801 | <0.0001 | Zhang et.al. 2021 |
| Five lncRNAs (AC069513.4, AC003092.1, CTC-205M6.2, RP11-507K2.3,U91328.21) | Bulk transcriptome | 0.68 | - | Shi et.al. 2016 |
| miR-21 or miR-106a | Bulk transcriptome | 0.865 or 0.819 | 0.004 or 0.000 | Tusong et.al. 2017 |

